# Evolutionary conserved protein motifs drive attachment of the plant nucleoskeleton at nuclear pores

**DOI:** 10.1101/2021.03.20.435662

**Authors:** Sarah Mermet, Maxime Voisin, Joris Mordier, Tristan Dubos, Sylvie Tutois, Pierre Tuffery, Célia Baroux, Kentaro Tamura, Aline V. Probst, Emmanuel Vanrobays, Christophe Tatout

**Affiliations:** iGReD, Université Clermont Auvergne, CNRS, INSERM, 63001 Clermont-Ferrand, France; Université de Paris, CNRS UMR 8251, INSERM ERL U1133, Paris, France.; Department of Plant and Microbial Biology, Zürich-Basel Plant Science Center, University of Zürich, Zürich, Switzerland; Department of Environmental and Life Sciences, University of Shizuoka, Shizuoka 422-8526, Japan

## Abstract

The nucleoskeleton forms a filamentous meshwork under the nuclear envelope and contributes to the regulation of nuclear morphology and gene expression. To understand how the Arabidopsis nucleoskeleton physically connects to the nuclear periphery, we investigated the nucleoskeleton protein KAKU4 and sought for functional regions responsible for its localization at the nuclear periphery. Computational predictions identified three evolutionary conserved peptide motifs within the N-terminal region of KAKU4. Functional analysis revealed that these motifs are required for homomerization of KAKU4, interaction with the nucleoskeleton proteins CROWDED NUCLEI (CRWN) and localization at the nuclear periphery. We find that similar protein motifs are present in NUP82 and NUP136, two plant specific nucleoporins from the Nuclear Pore Complex (NPC) basket. These conserved motifs allow the two nucleoporins to bind CRWN proteins, thus revealing a physical link between the nucleoskeleton and nuclear pores in plants. Finally, whilst NUP82, NUP136 and KAKU4 have a common evolutionary history predating non-vascular land plants, KAKU4 mainly localizes outside the NPC suggesting neofunctionalization of an ancient nucleoporin into a new nucleoskeleton component.

## INTRODUCTION

The nuclear envelope has emerged as one of the key structures of the eukaryotic cell. It separates the nucleus from the cytoplasm, creating two functional compartments with replication and transcription taking place inside the nucleus and translation in the cytoplasmic compartment (1).

The nuclear envelope is made of a double membrane that is however more than a simple barrier. The outer nuclear membrane is connected to the cytoskeleton and the inner nuclear membrane to the nucleoskeleton (1). Functional components such as the LInker of Nucleoskeleton and Cytoskeleton (LINC) complex made of Klarsicht/ANC-1/Syne Homology (KASH) and Sad1/UNC84 homology (SUN) domain proteins bridge the two nuclear membranes and propagate extranuclear signals across the nuclear envelope into the nucleus to ultimately influence gene expression (2, 3). The nuclear envelope also contains several hundreds of Nuclear Pore Complexes (NPCs) that regulate the trafficking of macromolecules inside and outside the nucleus, and which play significant roles in transcription regulation (4, 5). Finally, the nuclear membrane is also tightly linked with the nucleoskeleton, a layer of filamentous structures maintaining the nuclear morphology and tethering specific chromatin domains to the nuclear periphery (6, 7). There is increasing evidence that the nuclear envelope, NPCs and the nucleoskeleton are closely connected to perform their function in a coordinated manner, but these interactions remain poorly understood especially in plants.

The NPC is composed of several units named nucleoporins (NUP) assembled into three distinct regions: the cytoplasmic fibrils, a central scaffold region, which makes up the central channel and a NPC basket located at the nucleoplasmic side of the NPC (8). Twenty-two novel NUPs have been identified in *Arabidopsis thaliana* including plant-specific NUPs such as NUP1/136 (hereafter NUP136) (9). NUP136 localizes to the NPC basket, contains Phenylalanine-Glycine (FG) repeats that regulate NPC permeability (10) and has been proposed to be the functional homolog of the yeast Nup1p and animal NUP153 (11, 12). NUP136 interacts with NUP82, another plant specific NUP, and the two proteins share sequence similarities within their N-terminal part (13). NUP136 also interacts with THP1, a component of the TREX-2 mRNA export complex (14) and NUP136 loss of function induces accumulation of mRNA within the nucleus, consistent with its conserved function in mRNA export (9). Although there is so far no evidence for direct interaction of NUP136 with DNA, NUP136-GFP was shown to bind chromatin regions reminiscent to the B compartment previously revealed by Hi-C experiments, which tend to be gene-poor and enriched in repressed chromatin (15).

The plant nucleoskeleton has also a unique composition, which do not encode lamin homologs, but CROWDED NUCLEI (CRWN) proteins that only share long coiled-coil regions with their animal counterparts (16). Another component of the plant nucleoskeleton is KAKU4, also specific to plants, that interacts with CRWN1 and CRWN4 (17). Despite their different composition, the nucleoskeleton in animals and plants shares common functions such as maintenance of nuclear morphology (16–18). Interestingly, nuclear morphology is also influenced by NUP136, but not by other NUPs, suggesting a possible connection between NUP136 and the nucleoskeleton (19). In animals, lamins are known to interact with specific genomic regions and to create distinct chromatin folding domains called Lamin Associated Domains (LADs) (20). Plant LADs have been recently characterized and as for NUP136 associated regions, they correspond to the repressed B compartment, which includes also transposable elements that interact with CRWN1 (21). In addition, chromatin regions enriched in CRWN1 largely overlap with those binding NUP136 (21) raising the possibility of a possible dual contribution of NPC and nucleoskeleton to promote contact between repressed chromatin and the nuclear envelope.

Here we have chosen to investigate protein-protein interactions of KAKU4 as an entry point to better understand how the nucleoskeleton is tethered at the nuclear periphery. We identified short peptide motifs in Arabidopsis KAKU4 and investigated the function of three of these motifs. We show that they are required for interaction with CRWN proteins and for localization of KAKU4 at the nuclear periphery. Unexpectedly, we find that these motifs are conserved in NUP82 and NUP136 located in the NPC basket. The conserved peptide motifs in the two nucleoporins also mediate interaction with the CRWN protein family demonstrating that the plant nucleoskeleton is connected to the NPC through direct protein-protein interaction with the NPC basket. Given this functional and structural analogy to animals (22, 23), this suggest that similar functions in plants and animals are achieved by different proteins at the nuclear periphery. Finally, our phylogenetic analyses suggest that KAKU4 has diverged from an ancestral plant-specific nucleoporin, which had the ability to interact with the nucleoskeleton, to become itself a new component of the plant nucleoskeleton.

## MATERIALS AND METHODS

### Gene and Protein sequences

Orthologs of KAKU4, NUP136 and NUP82 were collected from 22 plant species that best represent the evolutionary history of the green lineage: *Arabidopsis thaliana* (A.th.), *Arabidopsis lyrata* (A.ly.), *Brassica rapa* (B.ra.), *Carica papaya* (C.pa.), *Glycine max* (G.ma.), *Theobroma cacao* (T.ca.), *Vitis vinifera* (V.vi.), *Populus trichocarpa* (P.tr.), *Prunus persica* (P.pe.), *Solanum lycopersicum* (S.ly.), *Oryza sativa* (O.sa.), *Zea mays* (Z.ma.), *Sorghum bicolor* (S.bi.), *Ananas comosus* (A.co.), *Musa acuminata* (M.ac.), *Amborella trichopoda* (A.tr.), *Picea abies* (P.ab.), *Pinus taeda* (P.ta.), *Marchantia polymorpha* (M.po.), *Physcomitrella patens* (P.pa.), *Oestrococcus lucimarinus* (O.lu.) and *Chlamydomonas reinhardtii* (C.re.). Most genomes of these plant species are available at Phytozome 7.0 (24) and Monocots or Dicots PLAZA 4.5 (25). The gymnosperms genomes were collected from conGenIE.org (Conifer Genome Integrative Explorer) (26). Gene and protein accession numbers used in this study are listed in **Supplemental Table S1**.

### Protein structure predictions

Protein structure of KAKU4, NUP82 and NUP136 were defined using Uniprot (27) and pfam (28) databases for domain predictions.

Secondary structures were predicted using PHYRE2, QUARK and PEP-FOLD3 (29–31). Intrinsically disordered regions (IDRs) were predicted using DEPICTER, DISOPRED3 and VSL2B (32–34) (**Supplemental Table S2**). VSL2B is the baseline predictor of VSL2 that is using only the amino-acid composition-based features that can be calculated directly from the protein sequence (34). DescribePROT database was used to collect pre-computed IDR predictions for 27,466 proteins of the *Arabidopsis thaliana* proteome including those predicted by VSL2B (35). VSL2B predictions were extracted for 31 NUPs, 5 Nucleoskeleton proteins and 5 inner nuclear membrane proteins (**Supplemental Table S3**). All schemes of protein structures were drawn to scale using the drawProteins R package (36).

### Phylogenetic studies

Phylogenetic tree was constructed using protein sequences collected from 52 species using NGPhylogeny.fr (37) with MAFFT 7.407 for multiple alignment, BMGE 1.12 for alignment curation and PhyML 3.1 for tree generation with 1000 bootstrap replicates. Tree was refined using the Interactive Tree Of Life (ITOL) (38). The amino-acid substitutions rates were computed as the sum of the branch lengths (*i.e.* sum of the amino-acid substitutions per site) of the phylogenetic tree described above and by considering separately KAKU4, NUP82 and NUP136 clades. Only the branch lengths corresponding to the 16 species shared by the three clades were considered.

### Domain and motif search in proteins

The orthologs of KAKU4, NUP82 and NUP136 were identified using the Arabidopsis proteins as a query by performing an all-against-all protein sequence similarity search using BLASTP with an E-value cut-off of 10^−10^. Ortholog occurrence in distant species was refined using sequences from *Amborella trichopoda* (basal angiosperm species) with the same approach. The MEME 5.1.1 (Multiple Em for Motif Elicitation) suite was used for *de novo* motif predictions (39). From a list of orthologous proteins, MEME was parameterized to define 10 motifs, each with a maximum length of 100 amino-acids. Protein sequences from each clade (KAKU4, NUP82 and NUP136) were submitted to MEME either individually, or in combination of 2 or 3 with the assumption that the three protein families contain common conserved regions. MAUVE (40) was used to define Locally Conserved Block (LCBs) between genomic regions of about 100kb each centered on the *KAKU4*, *NUP82* and *NUP136* genes.

Nonsynonymous (dN) and synonymous (dS) substitution rates were investigated to determine if selection or genetic drift occurs within the *KAKU4*, *NUP82* or *NUP136* gene families. To avoid underestimation of dS, the analysis was restricted to 9 close relative species from Brassicaceae (*Arabidopsis thaliana, Arabidopsis lyrata, Camelina sativa, Capsella rubella, Eutrema salsugineum, Brassica napus, Brassica oleraceae, Brassica rapa, Raphanus sativus*). The Codeml program of the Phylogenetic Analysis by Maximum Likelihood (PAML) package (v.4.4c) (41) was used with the variable *NSsites* to compare the observed dN/dS ratio for KAKU4, NUP82 and NUP136 phylogenetic tree with the null model M0 where all tree branches have the same dN/dS set to 1 (no selection). The M0 model was rejected if P-value < 0.05.

### Constructs and cloning

cDNAs of *CRWN1* (*AT1G67230*), *CRWN*2 (*AT1G13220*), *CRWN3* (*AT1G68790), CRWN4 (AT5G65770), KAKU4 (AT4G31430), NUP136 (At3g10650*) and *NUP82* (*AT5G20200*) were cloned in pDONR vectors using gene-specific primer sequences (**Supplemental Table S4**). Motifs were amplified from the corresponding cDNA using specific primers (**Supplemental Table S4**) and cloned in pDONR vectors, while the domain deletions were constructs from initial cDNA constructs using the site-directed mutagenesis kit QuikChange™ (Agilent Technologies) and adequate primers (**Supplemental Table S4**). All cloning procedures rely on Gateway technology®. After initial cloning, full-length coding sequences and other constructs were cloned into appropriate expression vectors for yeast-two hybrid assays (bait vector pDEST- GBKT7 or prey vector pDEST-GADT7), expression *in planta* (pGWB606-GFP, pGWB555-RFP) or BiFC (pB4cYGW, pB4nYGW, pB4GWnY and pB4GWcY). The list of all the plasmids used in this study can be found in **Supplemental Table S5** and **S6.**

### Yeast Two-Hybrid Assay

Yeast cultures were grown at 30°C on YPD or on selective SD media. *Saccharomyces cerevisiae* strains AH109 Gold and Y187 (Clontech, MATCHMAKER GAL4 Two-Hybrid System) were transformed according to the classical protocol (42) separately by the bait vector pDEST-GBKT7 or the prey vector pDEST-GADT7 (43) and grown on appropriate medium (SD-Trp or SD-Leu plates respectively). After mating on YPD and selection of diploids on SD-Leu-Trp medium, interactions were recorded on stringent medium SD-Leu-Trp-His-Ade. Empty pDEST-GBKT7 or pDEST-GADT7 vectors were used as negative controls.

### Transient expression of fusion proteins in tobacco

Vectors pGWB606 (GFP Nter), pGWB555 (mRFP Nter) (44) or pB4GWnY (Nter of YFP / nYFP Cter), pB4nYGW (Nter of YFP / nYFP Nter) pB4GWcY (Cter of YFP / cYFP Cter) and pB4cYGW (Cter of YFP / cYFP Nter) (45) were used for transient expression studies. Expression vectors were transformed in strain GV3101 of *Agrobacterium tumefaciens*. The p19 suppressor of gene silencing was used to enhance expression (46). Transient co-expression of the expression vectors and p19 constructs was performed by infiltration Agrobacterium cultures at OD^600^ of 0.1 and 0.05, respectively, into 5-6-week-old leaves of *Nicotiana benthamiana* as described before (47). Expression profiles are observed two days after agro-infiltration using confocal microscopy (LSM800, Zeiss).

### Image acquisition, storage and analysis

Fluorescence images were obtained using an inverted confocal laser-scanning microscope (LSM800; Carl Zeiss). The 488-nm line of a 40-mWAr/Kr laser, and the 544-nm line of a 1-mW He/Ne laser were used to excite GFP/YFP, and RFP, respectively. Images were acquired with 40x or 63x oil immersion objectives. All images are stored in an *in house* OMERO server and figures were produced using OMERO.figure (48). Images of **Supplementary Figure S3** were processed and rendered using Imaris (Bitplane AG).

## RESULTS

### The N-terminal part of KAKU4 is important for protein-protein interaction and its localization at the nuclear periphery

Previous experiments revealed that Arabidopsis KAKU4 and CRWN1 interact directly to form a complex together with CRWN4 (17). To gain further insight into the protein domains responsible for these interactions, we generated several constructs expressing either the full-length protein (amino-acids 1-574), the N-terminus (1–310) or the C-terminus (311–574) of KAKU4 (Materials and Methods and **Figure 1A**).

**Figure 1:**
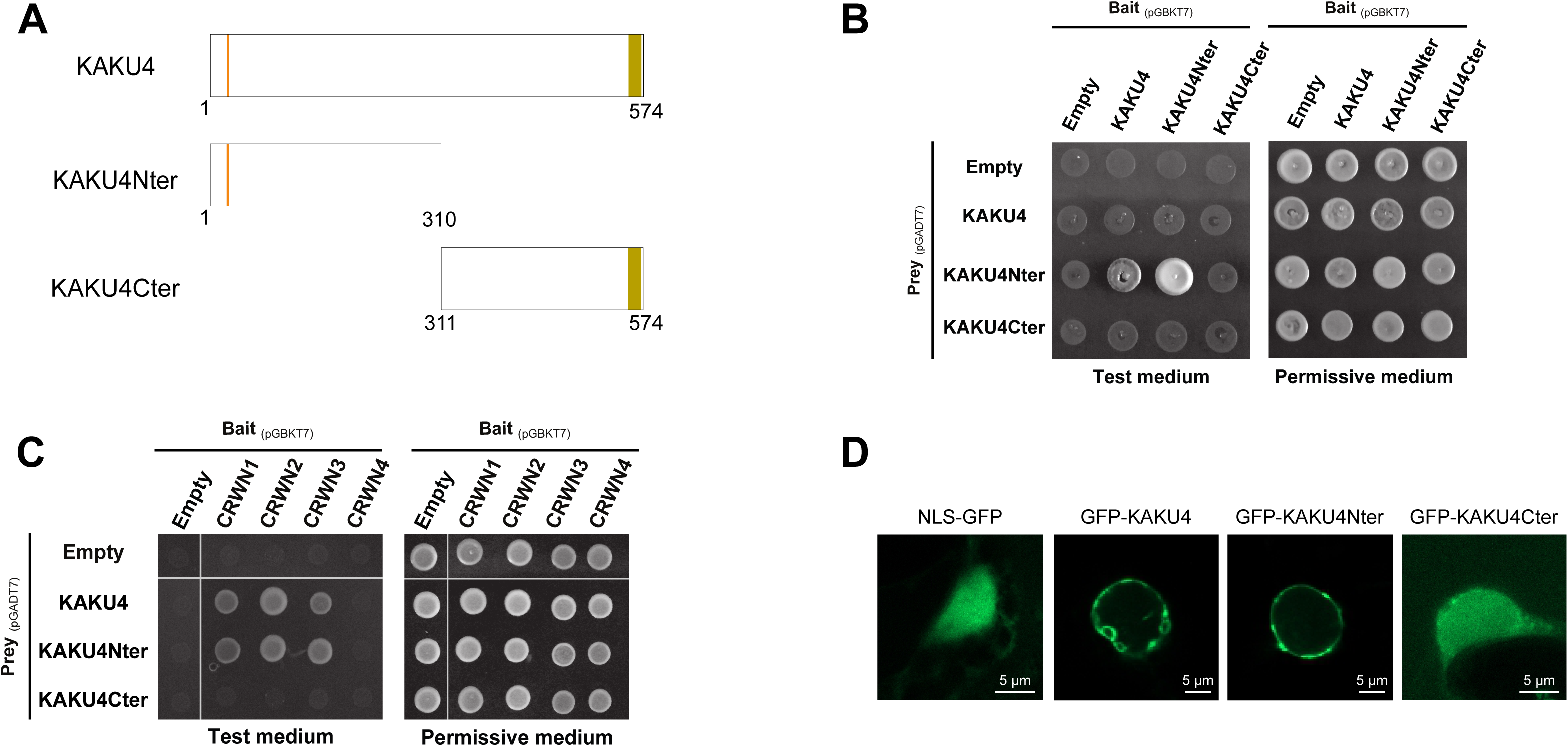
The N-terminal part of *Arabidopsis thaliana* KAKU4 is important for protein-protein interaction and localization at the nuclear periphery. **(A)** Protein organization of KAKU4 and deletion constructs used. **(B** and **C)** Yeast two-hybrid assay. Yeast strains were grown on Permissive (SD/–Leu/–Trp) or Test (SD/–Leu/– Trp/–Ade/–His) Media. Empty vectors pGBKT7 and pGADT7 are used respectively as bait and prey negative controls. **(B)** KAKU4 interaction with itself. **(C)** Interaction between CRWN1 to 4 and KAKU4 deletion variants. **(D)** KAKU4 localization at the nuclear periphery (green fluorescent signal). Confocal images of NLS-GFP, GFP- KAKU4 or deletion constructs transiently expressed in *Nicotiana benthamiana* leaf epidermal cells.

In combinatorial yeast-two-hybrid (Y2H) experiments the N-terminal region of KAKU4 is multifunctional and enables homomerization (**Figure 1B**) as well as interaction with CRWN1, CRWN2 and CRWN3 but not with CRWN4 (**Figure 1C**). Deletion of the N-terminal region completely abolishes interaction with the CRWN proteins (**Figure 1C**). To demonstrate the relevance of these results in plant cells, the same KAKU4 constructs were fused to GFP and transiently expressed in tobacco. We examined their nuclear localization to determine the contribution of N- and C-terminal regions of KAKU4 to its enrichment at the nuclear periphery and found that the N-terminal domain was required and sufficient for the localization at the nuclear periphery (**Figure 1D**).

The N-terminal region of KAKU4 thus plays a central role in allowing KAKU4 to interact with its protein partners and to ensure its localization to the nuclear periphery.

### The N-terminal region of KAKU4 contains conserved peptide motifs

To gain insight into the protein characteristics allowing for KAKU4 enrichment at the plant nuclear periphery, we first searched for protein domains that are evolutionary conserved in the KAKU4 protein sequence using referenced protein domains in pfam (28). This approach identified a predicted Nuclear Localization Signal (NLS) in the N-terminal part and a Glycine-Arginine rich (GAR or RGG/RG box (49)) region at the C-terminus (**Figure 2A**) but did not identify other conserved protein regions.

**Figure 2:**
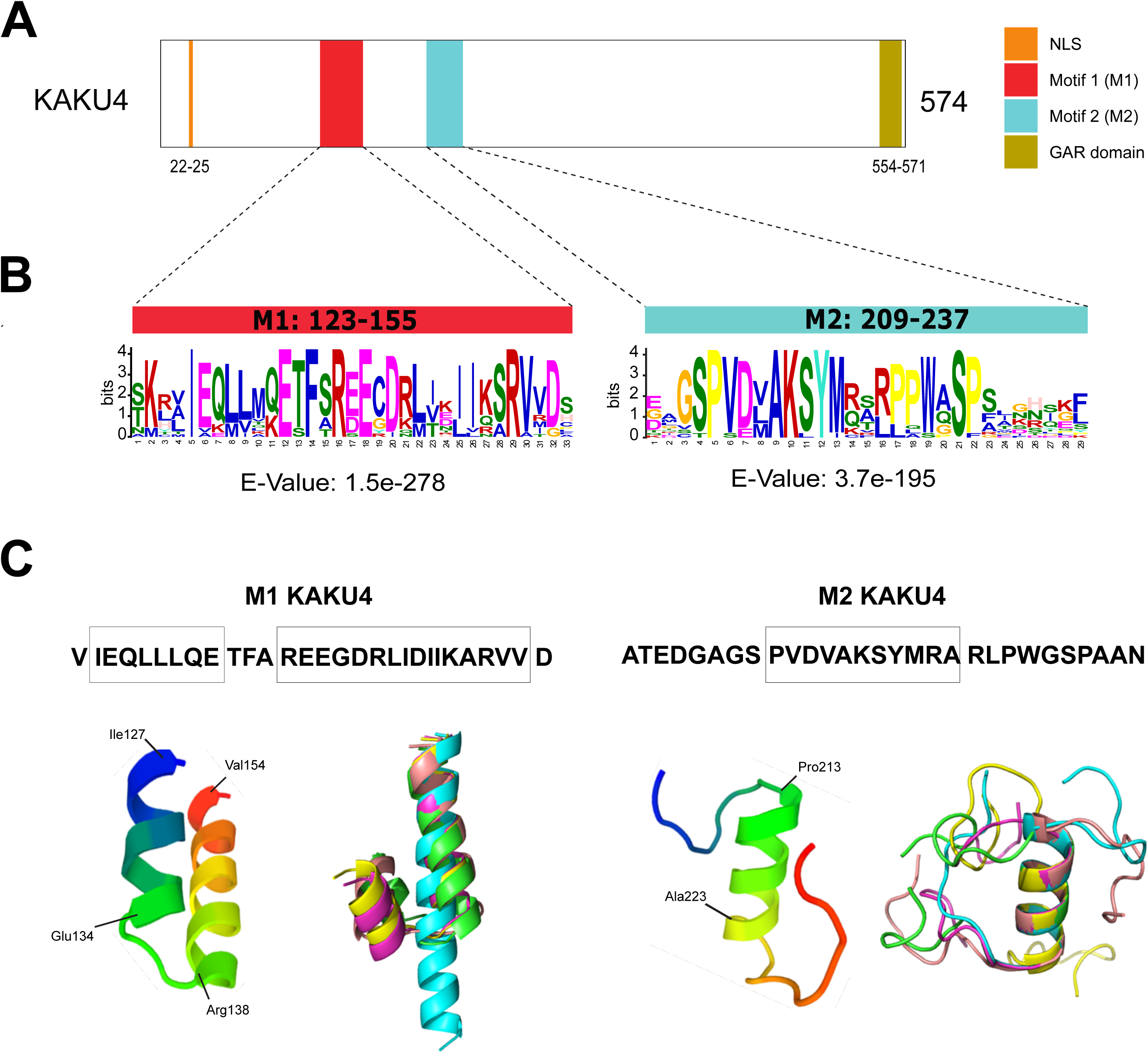
*A. thaliana* KAKU4 N-terminal region contains two conserved peptide motifs. **(A)** Domains and motifs organization of KAKU4 including a NLS, a glycine arginine rich domain (GAR domain) and two conserved motifs named M1 (123-155) and M2 (209-237) identified by MEME analysis using KAKU4 orthologs from 16 selected plant species (Materials and Methods). **(B)** Motifs identified by the MEME suite. Motifs correspond to the probability of each possible amino acid (letter code) at each position in the peptide sequence. The E-value significance of the two motifs from KAKU4 is indicated under each logo. **(C)** Ribbon diagram of the predicted folded structures using PEP-FOLD3. The peptide sequence used for prediction is indicated at the top of the figure and predicted helices are indicated as boxes. Best model (left) and superposition of the five best models (right) of M1 and M2 from *Arabidopsis thaliana* KAKU4 suggests a helical organization for M1 and M2. M1 helical region is possibly broken around residues 134-TFA-137.

Yet, the above database search is not powerful to detect small regions of less than 100 amino acids, which can potentially also contribute to protein interactions. Therefore, we deployed the MEME suite (39), to predict novel motifs conserved in 16 KAKU4 protein orthologs from basal to recent angiosperms (Materials and Methods). This approach revealed conserved regions in the N-terminal domain of KAKU4. The two most conserved motifs (M) were selected and numbered according to their E-value ranking (M1 and M2) (**Figure 2B**). Because M1 and M2 motifs are about 30 amino acids long, we use hereafter the generic term of peptide motifs instead of Short Linear Motifs (SLMs), which is usually referring to 5-15 amino-acid motifs. We then predicted folding structures of the N-terminal part of KAKU4 using the QUARK and PHYRE2 algorithms (29, 31), respectively, based on *ab initio* protein structure prediction and advanced homology search methods. Both algorithms predicted the presence of alpha helices overlapping with M1 and M2 (**Supplemental Figure S1**). The presence of alpha helices was confirmed by a third approach using the PEP- FOLD3 *de novo* structure modeling engine (30). The five best models predicted by PEP-FOLD3 for M1 and M2 sequences were superposed confirming the occurrence of alpha helices. More specifically, a helix region (I127-V154) possibly broken around residues 134-TFA-137 was defined for M1, and a shorter helix (P213-A223) was predicted for M2 (**Figure 2C**).

As alpha helices are known to be frequently involved in protein-protein interactions (50), our analyses suggest that the conserved motifs could account for the interaction properties observed for the N-terminal region of KAKU4.

### KAKU4 peptide motifs mediate protein-protein interactions

To test the function of the M1 and M2 motifs, two complementary types of constructs were designed. A first set expresses short, coding sequences restricted to M1 and/or M2 motifs while a second set expresses full-length KAKU4 versions from which M1 and/or M2 motifs were deleted (Materials and Methods) (**Figure 3A**).

**Figure 3:**
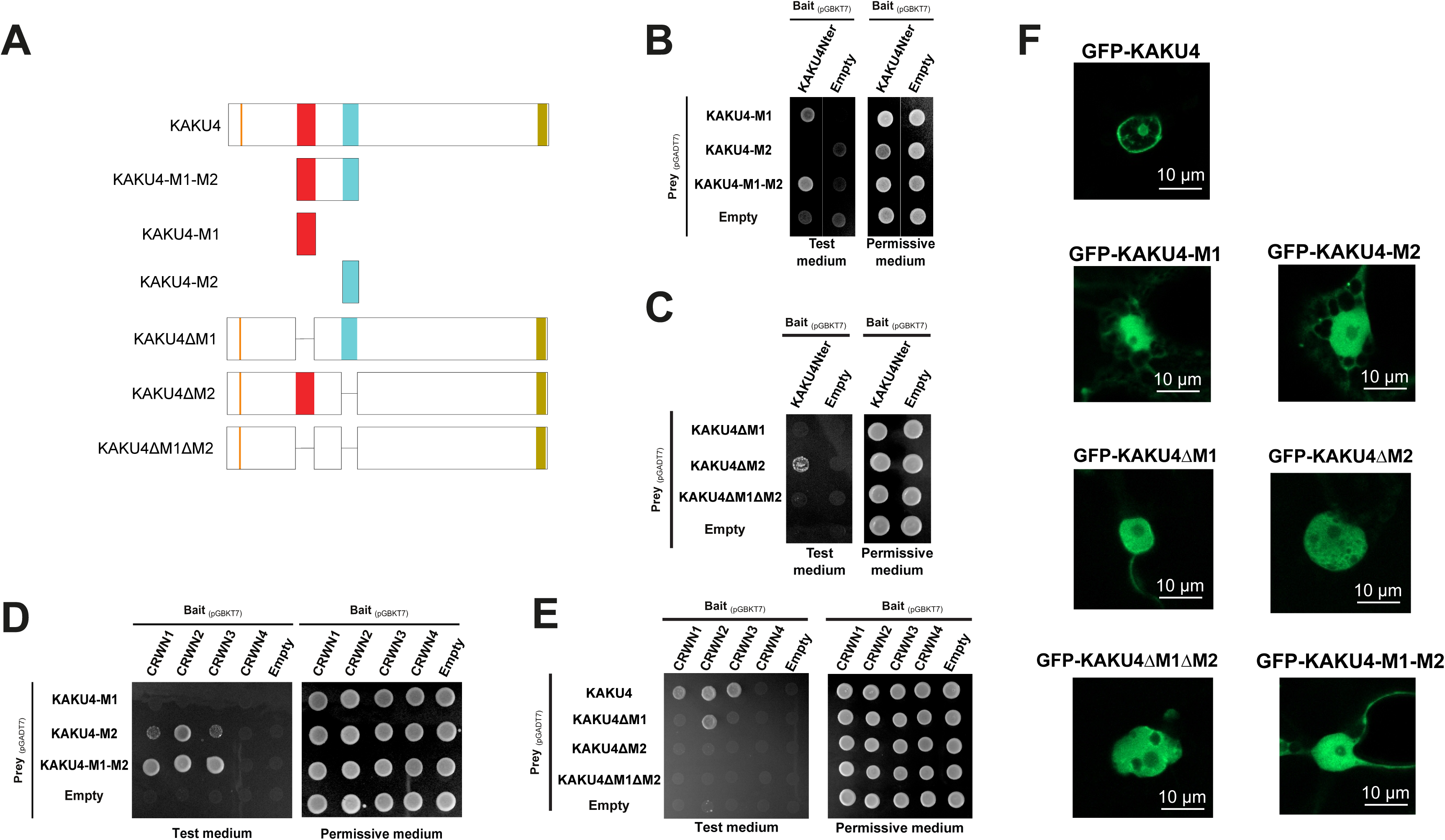
*A. thaliana* KAKU4 motifs M1 and M2 are necessary for protein-protein interactions and correct localization. **(A)** Schemes of the KAKU4 protein constructs containing either only M1 and / or M2 motifs or lacking these motifs. **(B-E)** Yeast two-hybrid assay on permissive (SD/–Leu/–Trp) or test (SD/–Leu/–Trp/–Ade/– His) media. Empty vector pGBKT7 and pGADT7 are used respectively as bait and prey negative controls. **(B and D)** Yeast-Two Hybrid assay testing interaction between motifs M1 and M2 from KAKU4 and the N-terminal region of KAKU4 **(B)** or the CRWN proteins **(D)**. **(C and E)** Yeast-Two Hybrid assay testing interactions between M1 and M2 deletions and the AtKAKU4 N-terminal region **(C)** or the CRWN proteins **(E). (F)** Confocal images of GFP-KAKU4 and various mutant constructs transiently expressed in *N. benthamiana* leaves.

We investigated by Y2H the ability of these synthetic proteins to interact with KAKU4 and CRWN proteins. The results showed that the M1 motif alone can interact with KAKU4, while a KAKU4 protein deleted of M1 loses this property, hence the M1 motif is required to promote interaction with KAKU4 (**Figure 3B-C**). In the same way, we were able to establish that M2 is necessary and sufficient for interactions with CRWN1, CRWN2 and CRWN3 proteins (**Figure 3D-E**). The localization of GFP-tagged KAKU4 variants was then analyzed by transient expression *in planta.* The absence of M1, M2 or both motifs impaired KAKU4 localization at the nuclear periphery (**Figure 3F**) suggesting that both homomerization of KAKU4 and interaction with CRWN proteins are required to localize KAKU4 at the nuclear periphery.

These results highlight the functional importance of the conserved M1 and M2 motifs in the N-terminal region of KAKU4. The motifs provide specificity in protein-protein interactions to form a complex with CRWN proteins and are necessary to localize *Arabidopsis* KAKU4 at the nuclear periphery.

### Conserved peptide motifs in KAKU4 are also found in two nucleoporins of the NPC basket

We then used the motifs identified by the MEME suite as query in a BLASTP search. (51) or using FIMO (52) to determine whether they are present in other proteins. Regions of similarity were indeed found in NUP82 and NUP136, two components of the NPC basket (9, 13) (**Figure 4A**). Motif prediction was then refined by performing a new MEME analysis for each of the three proteins or using combinations of orthologs from the two or three protein families. With this strategy, three common conserved motifs were identified (**Figure 4B**) among a set of 52 orthologs representing the *NUP82*, *NUP136* and *KAKU4* gene families (Materials and Methods and **Supplemental Table S1**): M1 and M2 previously identified in KAKU4 and a third motif M3 located N-terminally of the two other motifs (**Supplemental Figure S2**). The discovery of this new conserved motif prompted us to test whether M3 contributes to target Arabidopsis KAKU4 to the nuclear periphery. When testing the requirement of this motif for KAKU4 localization in transient expression assays, we found that M3 alone was not sufficient, but requires the presence of M1 and M2 to mediate KAKU4 localization at the nuclear periphery (**Figure 4C** and **4D**). We then wanted to compare in more detail the localization of NUP82, NUP136 and KAKU4 at the nuclear periphery. GFP-NUP136 shows a spotty pattern as expected for a NPC component (9), while GFP-NUP82 has a more regular pattern as for KAKU4-tRFP or forms large clusters at the nuclear periphery that might potentially be attributed to the overexpression in a heterologous system (**Supplemental Figure S3A**). Detailed image reconstruction using IMARIS shows that KAKU4-tRFP fusion proteins adopt a more uniform distribution at the nuclear periphery, interspersed by GFP-NUP136 foci (**Supplemental Figure S3B**). This, even though the three proteins share the three conserved motifs, KAKU4 show a different localization pattern than the two NUPs.

**Figure 4:**
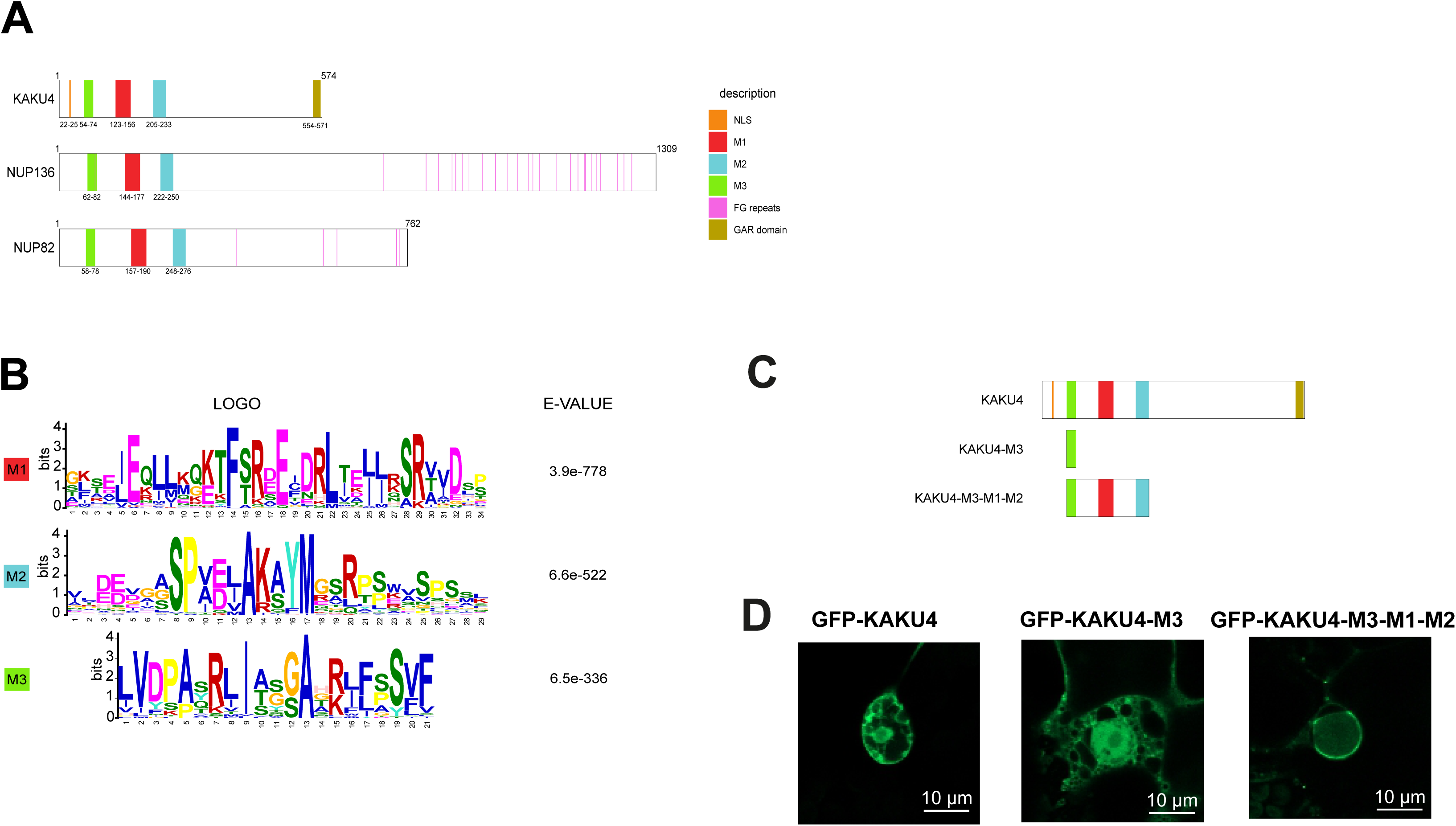
M1 and M2 motifs are conserved in nucleoporins NUP136 and NUP82. **(A)** Schematic representation of *A. thaliana* KAKU4, NUP136 and NUP82 including the three common motifs and other specific features (NLS, GAR domain, FG-repeats). **(B)** Logo for each motif generated by MEME (left) with their associated E-value (right) using orthologs of KAKU4 (16 orthologs), NUP82 (16 orthologs) and NUP136 (20 orthologs) from 20 plant species (Materials and Methods). **(C)** Schematic representation of AtKAKU4 protein constructs containing either only M1 alone or with the M2 motifs. **(D)** Confocal images of GFP-KAKU4 and GFP constructs comprising only M3 or the three motifs from KAKU4 transiently expressed in *N. benthamiana* leaves.

### NUP82 and NUP136 interact with nucleoskeleton proteins

To test whether the conserved motifs in the two NPC basket proteins would enable their interaction with the nucleoskeleton, we first performed Y2H assays with NUP82 and NUP136. While NUP136, when expressed as bait, auto-activated the reporter construct impeding to test its ability to homomerize, NUP82 can interact with itself (**Figure 5A** and **5B**). NUP82 and NUP136 weakly interact physically in Y2H and were unable to bind KAKU4 (**Figure 5A** and **5B**). However, Bimolecular Fluorescence Complementation (BiFC) assays confirmed that the two NUPs interact with each other as previously shown (13) and further revealed that NUP82 and NUP136 can also interact with KAKU4 *in planta* (**Figure 5C** and **Supplementary Table S7**). Similarly to KAKU4, the two NUPs interact directly with CRWN1, CRWN2 and CRWN3 proteins but not CRWN4 in Y2H assays although interaction was weaker for CRWN1 than for CRWN2 and CRWN3 (**Figure 5D**). Interaction between NUP82 and CRWN2 was confirmed *in planta* by BiFC (**Figure 5C**). Together with the Y2H assay, this suggests that the NPC basket connects nuclear pores with the nucleoskeleton.

**Figure 5:**
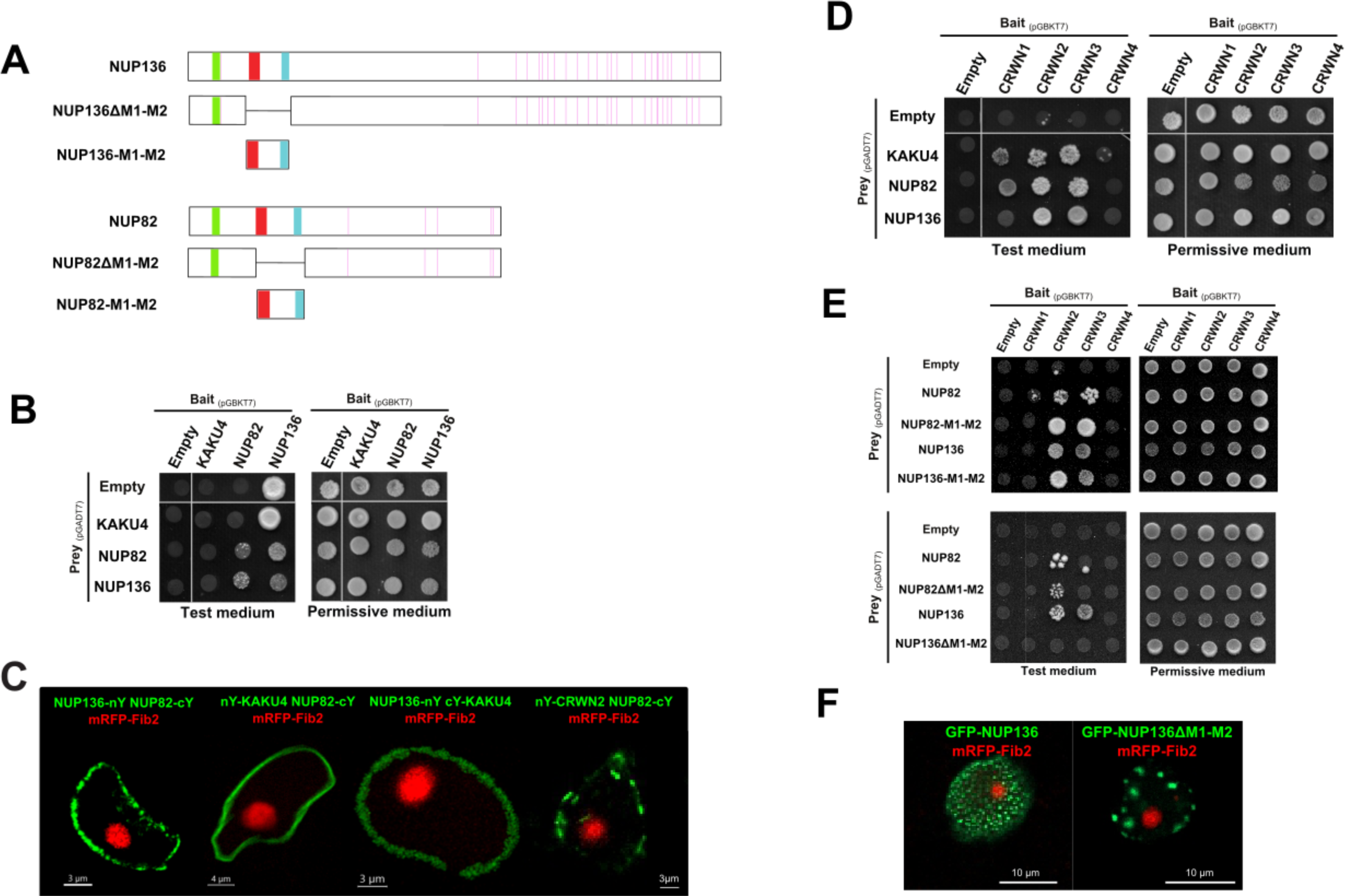
NUP82 and NUP136 interact with CRWN proteins via their M1 and M2 motifs. **(A)** Schematic representation of NUP136 and NUP82 and mutant constructs deleted for M1 and M2 motifs or expressing both motifs. **(B)** Yeast-Two Hybrid assay testing the interaction between KAKU4, NUP136 and NUP82 proteins. **(C)** BiFC assay in *N. benthamiana* leaves. Fibrillarin-RFP (red fluorescent signal) was co-expressed as a positive marker for transformed cells. nYFP and cYFP are the N-terminal and C-terminal half of YFP respectively (yellow fluorescent protein). **(D)** Yeast-Two Hybrid assay testing for interaction with CRWN proteins. **(E)** Yeast-Two Hybrid assay to test the contribution of M1, M2 motifs of NUP136 and NUP82 to the interaction with CRWN proteins. Empty bait and prey vector are used as negative control. **(F)** Confocal images of GFP-NUP136 and GFP-NUP136ΔM1-M2 constructs co-transfected with mRFP-Fibrillarin in *N. benthamiana* leaves.

NUP expressing constructs lacking the M1 and M2 motifs or containing only the two motifs were then tested for their ability to interact with CRWN proteins. In Y2H assays, M1 and M2 of NUP82 or NUP136 are sufficient to induce interactions with the CRWNs proteins (**Figure 5A** and **5E**). As shown in **Figure 5E**, stronger interactions were observed with CRWN2 and CRWN3 and deletion of the two motifs completely impaired these interactions for NUP136. For NUP82, residual interaction subsisted suggesting that NUP82 makes contacts with CRWN proteins outside the two motifs. When transiently expressed in tobacco as GFP fusion, removal of M1 and M2 did not affect the localization of GFP-NUP82 while the regular profile observed for GFP-NUP136 was lost and was leading to the formation of aggregates at the nuclear periphery (**Figure 5F** and **Supplementary Figure S4A** and **S4B**).

Taken together, these results confirm the contribution of the newly discovered conserved motifs M1 and M2 to protein-protein interaction and nuclear localization. They also demonstrate the existence of a protein complex between KAKU4, NUP82, NUP136 and the CRWN proteins at the periphery of the plant nucleus.

### KAKU4, NUP82 and NUP136 proteins contain intrinsically disordered regions

Short peptide motifs are often found in proteins with intrinsically disordered regions (IDR) that do not autonomously fold into a defined tertiary structure (53). IDRs are defined as regions lacking bulky hydrophobic amino acids and for this reason, they cannot form well-organized hydrophobic cores usually found in structured domains. However, when protein binding is initiated at the short peptide motifs, IDRs can undergo a disorder-to-order transition (54). IDRs can be computationally predicted based on amino-acid composition or by learning from sequence profiles surrounding residues unresolved (*i.e* unstructured) by crystallography (54). Many disorder predictors are available (55), here we used VSL2B an efficient predictor for proteins containing both structured and disordered regions (34), the meta predictors DISOPRED3 (33) and DEPICTER (32). KAKU4, NUP82 and NUP136 are predicted to contain several IDRs with values obtained for the full-length protein by three predictors ranging from 57.6% (VSL2, KAKU4) up to 96.5% (DEPICTER, NUP136) (**Figure 6A** and **Supplemental Figure S5A**). Examination of the N-terminal part of KAKU4, NUP82 and NUP136 (amino-acids 1 to 300) indicated several IDRs interspersed with structured regions overlapping the M1 and M2 peptide motifs (**Figure 6B**). This prompted us to investigate more proteins located at the NPC and its vicinity. For that purpose, we then took advantage of the DescribePROT database that provides access to pre-computed predictions from VSL2B for the *Arabidopsis thaliana* proteome (35). For the whole proteome, VSL2B predicts a median value of 36% of IDRs consistent with previous predictions (56). The level of IDRs among the NUPs is overall not exceeding the proteome average with a median value of 27.6% (among 31 NUPs, **Supplemental Table S3**). However, the NUPs localizing at the NPC basket such as NUP50a, NUP50b, NUP82, NUP136 and NUA all contain a higher proportion of disordered regions with a median value of 72.5% (**Figure 6C, 6D** and **Supplemental Figure S5B**). Such an enrichment in disordered regions was expected for NUP50a, NUP50b and NUP136 as they belong to FG-NUPs known to be intrinsically disordered proteins (57) but this enrichment was also observed for NUP82 and NUA as well as for CRWNs and KAKU4, a feature which was overlooked so far in these proteins (**Supplemental Figure S5B**).

**Figure 6:**
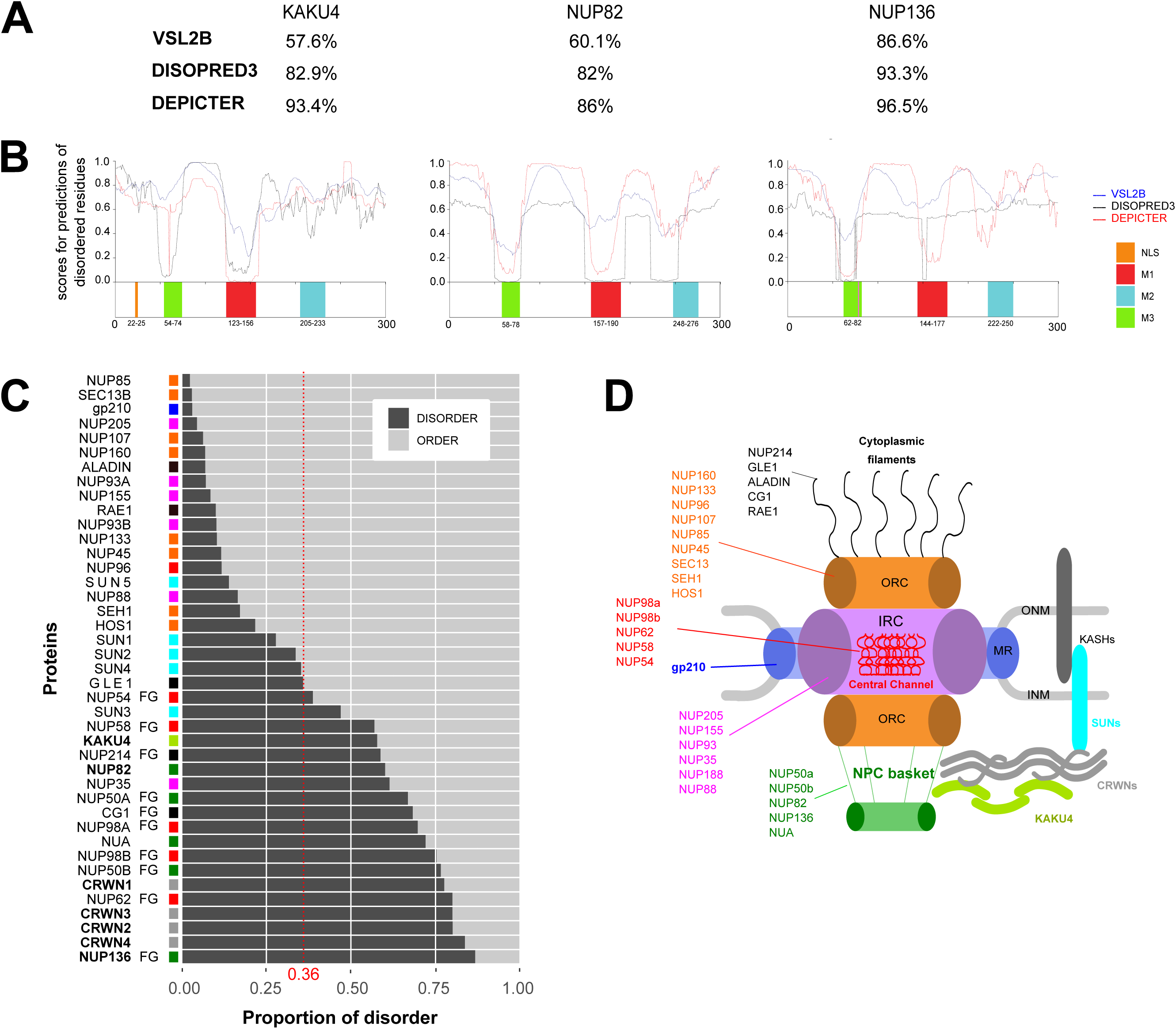
Predictions of intrinsically disordered regions in KAKU4, NUP136 and NUP82 orthologs. **(A)** Proportion of intrinsically disordered residues within full-length Arabidopsis KAKU4, NUP82 and NUP136 proteins predicted by VSL2B, DISOPRED3 and DEPICTER. **(B)** Profiles of intrinsically disordered residues within the N-terminal region (1-300) of KAKU4, NUP82 and NUP136. Schematic representations of the three N-terminal parts of the proteins are indicated below with M3, M1 and M2 peptide motifs indicated in green, red and blue respectively. **(C)** Proportion of intrinsically disordered regions (dark grey) and ordered regions (light grey) in NUPs, CRWNs, KAKU4 and SUN proteins from *A. thaliana*. The median value of disorder residues of the Arabidopsis proteome is indicated as a red dashed line. FG-NUPs and NPC sub-structures are indicated in a color code according to figure 6D on the left hand of the diagram. **(D)** Representation of the NPC structure with its main substructures and listing the different NUPs for each substructure in different colors. A schematic representation of KASH, SUN, CRWN and KAKU4 connections is given on the right of the figure.

These predictions suggest that proteins associated with the NPC basket and the nucleoskeleton tend to be structurally disordered. More specifically, KAKU4, NUP82, and NUP136 may not fold in stable secondary structures in their monomeric form except within their N-terminal part that contains the conserved M1, M2 and M3 motifs, which may dock the proteins within multimeric complexes.

### KAKU4 shares a common evolutionary origin with NUP82 and NUP136

The structural similarities observed between KAKU4, NUP82 and NUP136 led us to investigate whether they have a common evolutionary origin.

First, the presence of the three proteins was sought in a set of representative species of the green lineage for which the genome was sequenced (Materials and Methods). We observed that NUP136 is present in non-vascular land plants *Marcantiophyta*, conserved in *Bryophyta* and in all Angiosperms, which suggests that according to TimeTree (58), NUP136 appeared about 593 Million years ago (Mya). KAKU4 and NUP82 emerged more recently. They are both encoded by the basal Angiosperm, *Amborella trichopoda* genome, but the analysis including plant species before and after basal Angiosperm diversification (59), did not allow to resolve the exact timing of KAKU4 and NUP82 emergence. This identified *NUP136* as the most ancestral gene family with one to four paralogs in each of the analyzed species (**Figure 7A**).

**Figure 7:**
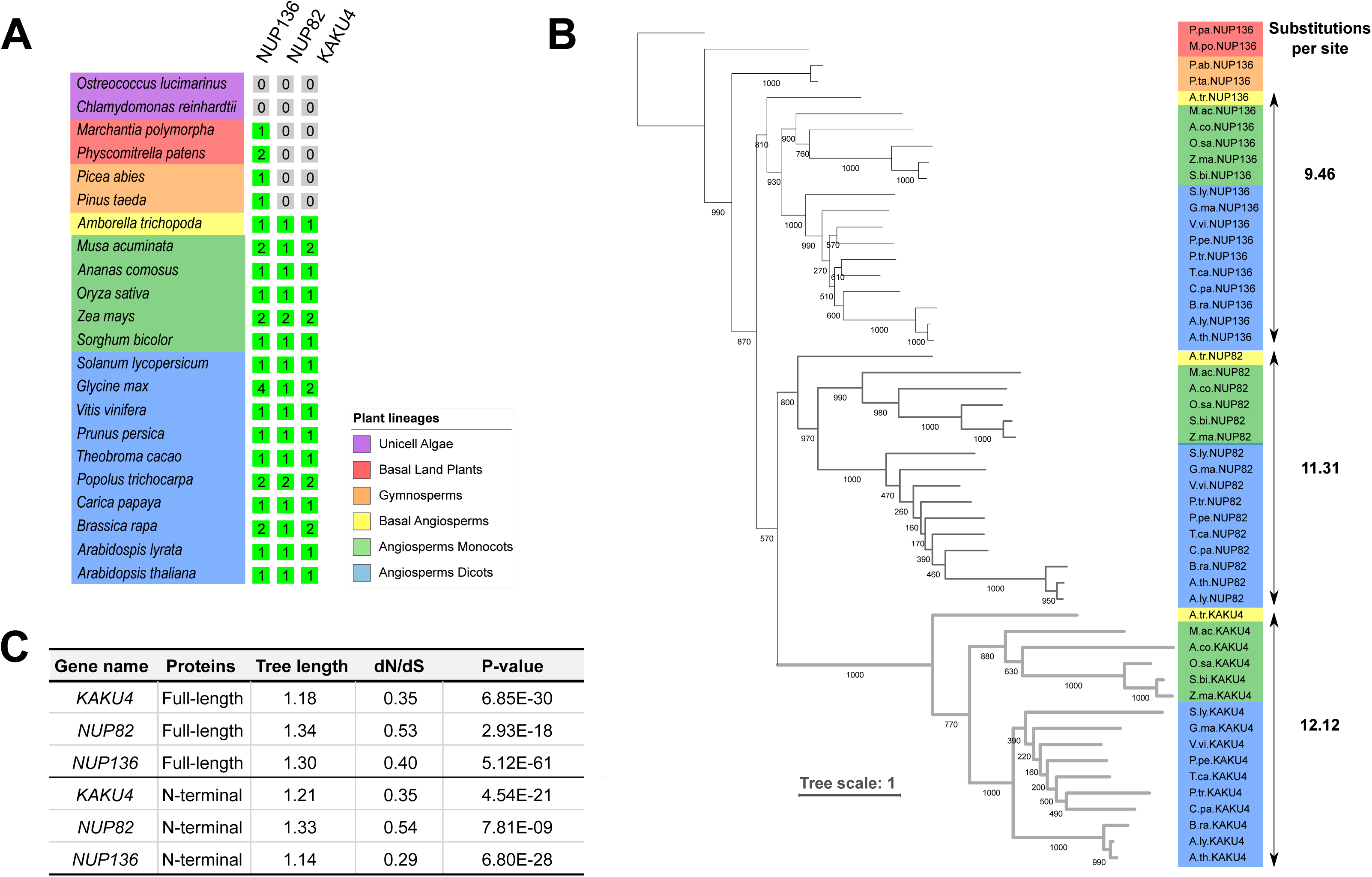
Phylogenetic relationships among KAKU4, NUP82 and NUP136. **(A)** Presence or absence of KAKU4, NUP82 and NUP136 proteins in the green lineage. The 22 plant species used in this study (left) and numbers of paralogs per species (right) for *NUP136*, *NUP82* and *KAKU4* genes are indicated. Selected species from top to bottom are from Unicellular algae (purple), Basal land plants (red), Gymnosperms (orange), Basal Angiosperms (yellow), Monocotyledons (green), and Dicotyledons (blue). **(B)** Maximum likelihood tree constructed from an alignment of amino-acid sequences of all orthologs from A). Bootstrap values are indicated on each branch. The substitutions per site are given at the right of the tree for each clade. The vertical arrows define the 16 species selected to estimate the sum of tree length for each clade. Clades are distinguished by thin dark branches (NUP136), thick dark grey (NUP82) and thick light grey (KAKU4). (**C)** Results of the test to determine whether the evolution of KAKU4, NUP82 and NUP136 is random (genetic drift) or under selection using Codeml NSsites from the PAML package. P-values are given for the full-length or N-terminal part of KAKU4, NUP82 and NUP136. Genetic drift hypothesis is rejected if P-value<0.5.

If *NUP136* is considered as the parental gene, the appearance of *KAKU4* and *NUP82* could have occurred after a gene or segmental duplication of a chromosomal segment including the *NUP136* gene or following an ancient genome-wide duplication (60). In order to search for traces of such an event, genomic context of 100kb flanking the three loci were investigated using multiple genome alignments (MAUVE) (40). In the monocot species *Zea mays*, Locally Collinear Blocks (LCBs), which are conserved segments that appear to be internally free from genome rearrangements, were identified at the immediate vicinity of *KAKU4, NUP82* and *NUP136* loci suggesting that the three chromosomal regions have a common origin (**Supplemental Figure S6A**). LCBs were also detected for *A. trichopoda* and *A. thaliana* (eudicot, modern angiosperm) though of much smaller size and more dispersed than in *Zea mays* (**Supplemental Figure S6B-C**). This strongly suggests that in angiosperms *KAKU4* and *NUP82* emerged from segmental or whole genome duplications of regions containing *NUP136*.

To illustrate the relationship between the three gene families, we then constructed a phylogenetic tree including all homologs (**Figure 7B**). The tree includes 52 proteins organized in three connected clades with different branch lengths suggesting different evolutionary rates for each clade. This was quantified by estimating the sum of the number of amino-acid substitutions per site for each clade (**Figure 7B**). The lowest value was observed for the NUP136 clade (9.4 substitutions per site), and the highest for the KAKU4 clade (12.1 substitutions per site). These values may argue in favor of a higher conservation of NUP136 protein sequence, which may reflect the preservation of its original function within the NPC basket while KAKU4 diverged faster. We then used the codon substitution modeling to determine if the observed divergence occurs randomly or if selection apply at nonsynonymous changes (41). For that purpose, we used Codeml *NSsites* on the whole cDNA or only the N-terminus and observed that selection indeed occurs in the three gene families (**Figure 7C**).

Collectively, our evolutionary analyses suggest a scenario of functional diversification of KAKU4, NUP82 and NUP136, whereas we propose *NUP136* as the parental gene ancestor with a biological function at the NPC, while NUP82 and KAKU4 diversified in protein composition and likely biological function.

## DISCUSSION

Our initial investigation of the KAKU4 protein led us to the discovery of three conserved motifs within its N-terminal region called M1, M2 and M3. These motifs range from 20 to 30 amino acids according to the MEME prediction or even smaller if only residues involved in secondary structures predicted by PEP-FOLD3 are considered. Here we could show that these motifs share at least two biological functions including protein-protein interaction (M1, M2) and targeting KAKU4 at the nuclear periphery (M1, M2 and M3). M1 and M2 are further involved in different types of interactions, including homomerization (M1) and multimeric complex formation with the CRWN proteins (M2). These motifs are not only present in KAKU4 homologs but also in the N-terminal part of NUP82 and NUP136, two nucleoporins of the NPC basket. Short peptide motifs have been described in NUPs from other species such as NUP93/Nic96, NUP98/Nup145N and NUP35/Nup53 (human/yeast, respectively) where these motifs mediate key interactions needed for the assembly of the inner ring complex of the nuclear pore basket with NUP188/Nup188, NUP205/Nup192 and NUP155/Nup170, respectively (8, 61). Interestingly, although the M1, M2 and M3 motifs of the plant NUP82, NUP136 and KAKU4 do not share sequence similarities with the motifs in yeast and human NUPs, they are functionally similar in promoting protein-protein interactions. Extending the approach described in this study to all plant NUPs may help unravel the structural basis for protein assembly at the plant NPC.

We also observed that NUP82, NUP136 and KAKU4 proteins contain large IDRs, which are interspersed with structured regions containing the identified peptide motifs in the N-terminal tail. IDRs may function as flexible linkers favoring the exposure of the motifs to facilitate interactions with other proteins while also providing a conformational separation of these protein-docking regions compatible with the formation of a multimeric complex. This scenario is reminiscent of the entropic chain- class of IDRs found for example in the 70 kDa subunit of replication protein A (RPA70), a eukaryotic protein that is essential for replication, recombination, and DNA repair (54). RPA70 contains an intrinsically unstructured linker region tethering two structured domains located at the center of the protein to a third structured DNA binding domain located at its N-terminus, called DBD F. The flexibility of the RPA70 allows DBD F to interact with DNA or with other proteins and mutations in the linker region of yeast RPA70 have been shown to impair its response to UV-induced DNA damage (62). We hypothesize that for NUP82, NUP136 and KAKU4, conformational transitions between fully disordered and partially structured folding states triggered by motif-mediated protein-protein interaction, may allow alternating between NPC and nucleoskeleton-specific functions. The occurrence of motifs with functional significance within disordered proteins may be more widespread in plants. While it is well documented in other species (54), few examples are known in plants (63), but efforts to identify them are starting to raise awareness on the role played by these proteins notably in nuclear functions (this study and (64)).

Here we revealed that KAKU4 shares the peptide motifs M1, M2 and M3 with the nucleoporins NUP82 and NUP136, where these motifs are involved in direct interactions with nucleoskeleton components CRWN and KAKU4. Possibly, in the absence of transmembrane domains in CRWN and KAKU4, these protein-protein interactions with nucleoporins may help to anchor the nucleoskeleton at the nuclear envelope. Besides nuclear pores, other anchoring points of the nucleoskeleton may exist, such as the inner nuclear envelope proteins, SUN. SUN1 was shown to bind CRWN1 (65) and the C-terminal fragment of CRWN1 (799–1132) interacts with SUN1 and SUN2 (66). Furthermore, using subtractive proteomics between NE enriched and NE depleted extracts as well as proximity-labeling proteomics, CRWNs, KAKU4 and SUN1 were found to be in close proximity at the nuclear envelope (67). As SUNs are sitting within the inner nuclear membrane through their transmembrane domains, it can be expected that SUNs contribute to anchor the nucleoskeleton at the nuclear periphery. Following this line of thought, KAKU4 may localize to the nuclear periphery not only through its interaction with nucleoporins and SUNs but also through its interaction with CRWN1, CRWN2 and CRWN3 and that its M2 motif would be central for these interactions. Experimentally, it remains challenging to test what drives KAKU4 localization at the nuclear periphery as the CRWN proteins display redundant functions and because depletion of several CRWN proteins strongly impairs plant viability (68).

An alternative, but not exclusive, role for the interactions between nucleoporins and the nucleoskeleton may be to achieve a regular distribution of NPCs within the nuclear envelope. NUP136 was proposed to be the functional counterpart of the human nucleoporin NUP153 (11). NUP153 interacts with lamin A and B and this interaction is required to recruit and maintain NUP153 at the NPC (22, 69). In mouse embryonic fibroblasts, removal of all lamins leads to clustering of the NPCs, which can be rescued by re-expression of either A or B-type lamins (70). Consistent with this, Arabidopsis NUP136 does not localize properly at the nuclear envelope when interactions with CRWN proteins are impaired through M1, M2 motif deletion, an observation in line with a model where the nucleoskeleton regulates NPC distribution at the nuclear membrane. Furthermore, interactions between the NPCs and the nucleoskeleton create a physical connection, possibly enabling a mechanical continuum from the nuclear envelope to the nuclear interior.

The role of mechanical transduction in regulating gene expression is increasingly discussed (71, 72). Since CRWN proteins also interact with genomic regions (15), and chromatin modifiers influencing gene expression (73, 74), complexes associating CRWN proteins with the nucleoskeleton and/or the NPC may provide a molecular mechanism to the hypothesized rheostat function of the nuclear envelope (75). Beyond the mechanotransduction hypothesis, the molecular connection between the nucleoskeleton and the NPC may provide a mechanism to induce or enhance gene expression upon external stimuli. Indeed, rapid responses involving one or several of the NUPs, KAKU4 and CRWN proteins have been reported during hyperosmotic stress (71), in response to copper in the medium (76) or upon biotic stress (77). Consistently, several examples suggest that gene expression in plants is influenced by the positioning of genes in respect to the nuclear periphery (76, 78–80). As such, the nucleoskeleton may play a central role by recruiting specific chromatin domains at the nuclear periphery (21, 74). NPCs are also known to participate in stress response (82). More specifically NUP82 and NUP136 control the expression of defense-related genes. This can be achieved either by direct interaction between NUPs and target genes or by modulating the nuclear import of specific regulators (5, 13). Thus, the physical link between NPC basket and the nucleoskeleton may contribute to rapid transitions between an inactive (*i.e.* near the nucleoskeleton) and active states (*i.e.* near the NPCs) in response to stress, as suggested during the formation of gene loops in yeast and animals (82). Investigation of sub-nuclear localization of NUP82, NUP136 and CRWN may help to determine in the future if chromatin domains with different transcription activity can be found at the NPC- nucleoskeleton boundary.

Finally, our phylogenetic analysis traced NUP136 back to the basal land plants and suggested that NUP82 and KAKU4 emerged from an ancestral duplication. One interesting hypothesis could be that two subsequent polyploidy events termed ξ (∼ 319 Mya) and ε (∼192 Mya), which according to TimeTree occurred respectively soon before (∼355 Mya) and after Angiosperm’s divergence (∼170 Mya) were responsible for these new duplication events (58). During evolution KAKU4 diverged into a new component of the nucleoskeleton, but still conserved some ability to interact with the NPC. This work highlights the tight relationship between the nuclear envelope, nuclear pores and nucleoskeleton. As part of this network the KAKU4 protein represents an intriguing case of neofunctionalization of an ancestral nucleoporin into a new component of the nucleoskeleton.

## ACKNOWLEDGMENTS

All pictures were acquired on the CLIC microscopy facility (CLermont Imagerie Confocale). CTa would like to thank Lukasz Kurgan for his advices on the use of DescribeProt and Antoine Molaro for critical reading.

---

**Supplemental Table S1: Genes and proteins IDs.** Gene IDs well collected from NCBI ref_seq, PLAZA (Monocots and Dicots 4.5) or Congenie.org ID and protein IDs from Uniprot database. IDs were collected from 24 species from the green lineage. Species are listed as full names or abbreviations as used in fasta files or phylogenetic trees. IDs are also grouped according to whether they belong to one of the KAKU4, NUP82 and NUP136 protein family.

**Supplemental Table S2: Intrinsically disorder prediction by VSL2, DEPICTER and DISOPRED3.** Score for prediction of disordered residues ranging from 0 (order) to 1 (disorder) are predicted for each amino-acid of KAKU4, NUP82 and NUP136 using VSL2, DEPICTER and DISOPRED3. Profiles using full-length and N-terminal protein sequences are depicted respectively in Supplemental Figure S4 in Figure 6A.

**Supplemental Table S3: Intrinsically disorder prediction by VSL2 for NUPs, nucleoskeleton and SUN domain proteins.** Protein and Gene names, Uniprot IDs is given and classified into NUPs, NSK (nucleoskeleton) and Inner Nuclear Membrane (INM) proteins. FG-NUPs and NUPs from the NPC basket are indicated. The number of ordered and disordered amino-acids (AA) were defined using precomputed predictions available at DescribeProt server (Materials and Methods) and then used to compute the proportion of disorder for each protein.

**Supplemental Table S4: Primer sequences used in this study.**

**Supplemental Table S5: Plasmid backbones used in this study.**

**Supplemental Table S6: Overview of constructs used in this study.**

**Supplemental Table S7: Bimolecular fluorescence complementation (BiFC) assay in *Nicotiana benthamiana* leaves.** Fibrillarin-RFP was co-expressed as a marker for visualizing nuclei in the transformed cells. nYFP and cYFP are the N-terminal and C-terminal half of YFP respectively. “+” positive interaction, “+/-” weak interaction; “-” absence of recorded interaction, “NT” not tested.

**Supplemental Figure S1:**
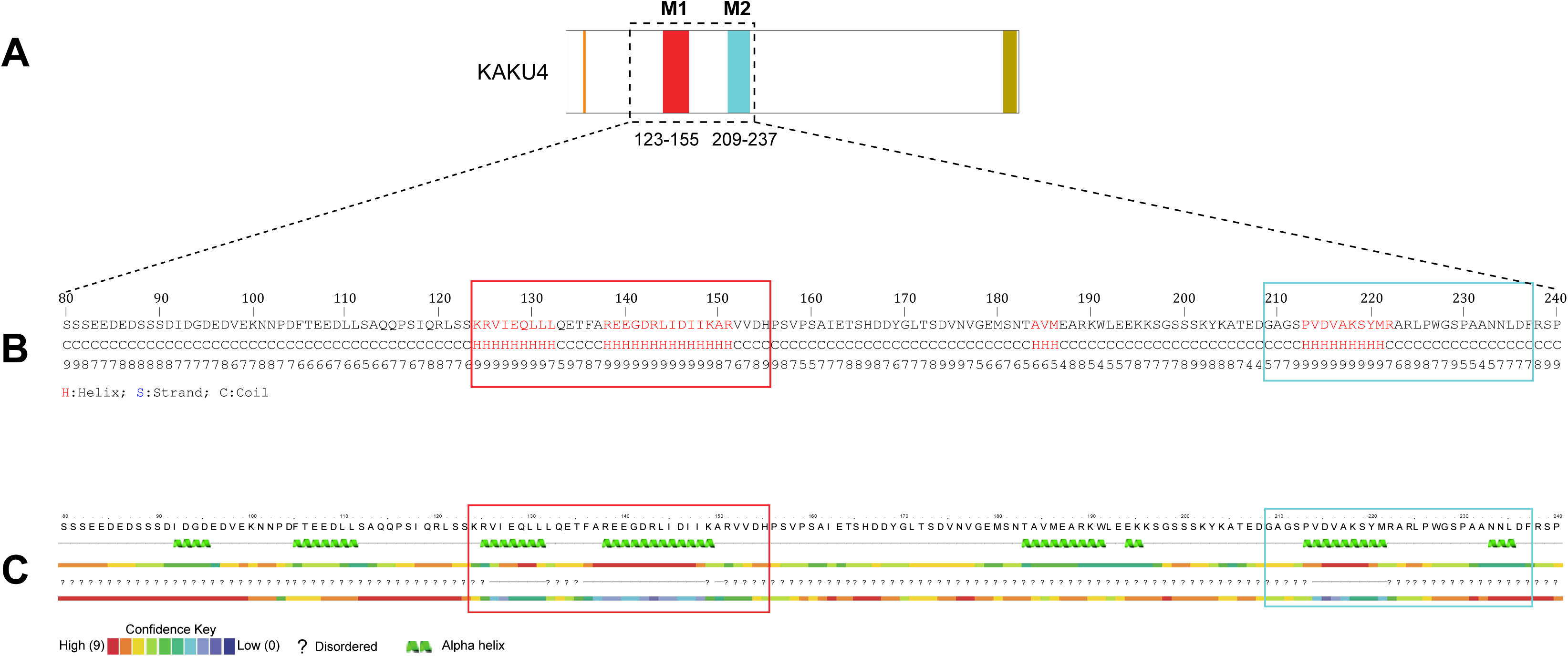
Structure predictions of *Arabidopsis thaliana* KAKU4 with QUARK, and PHYRE2 predictors. QUARK and PHYRE2 tools were used to investigate the possible secondary structure of the KAKU4 N-terminal region (1-240). **(A)** KAKU4 scheme with M1 and M2 peptide motifs indicated in red and blue respectively. **(B)** QUARK (Xu & Zhang, 2012) is an *ab initio* (model-free) method that calculates energy scores based on the force field exerted by the atoms constituting the amino-acids of the protein sequence. Confidence scores of the predicted secondary structure are indicated below the amino-acid sequence. **(C)** PHYRE2 (Kelley et al 2015) bases its predictions on models built from homologous proteins with a known structure. The M1 and M2 motifs detected by MEME are indicated respectively as red and blue boxes within the QUARK and PHYRE2 predictions.

**Supplemental Figure S2:**
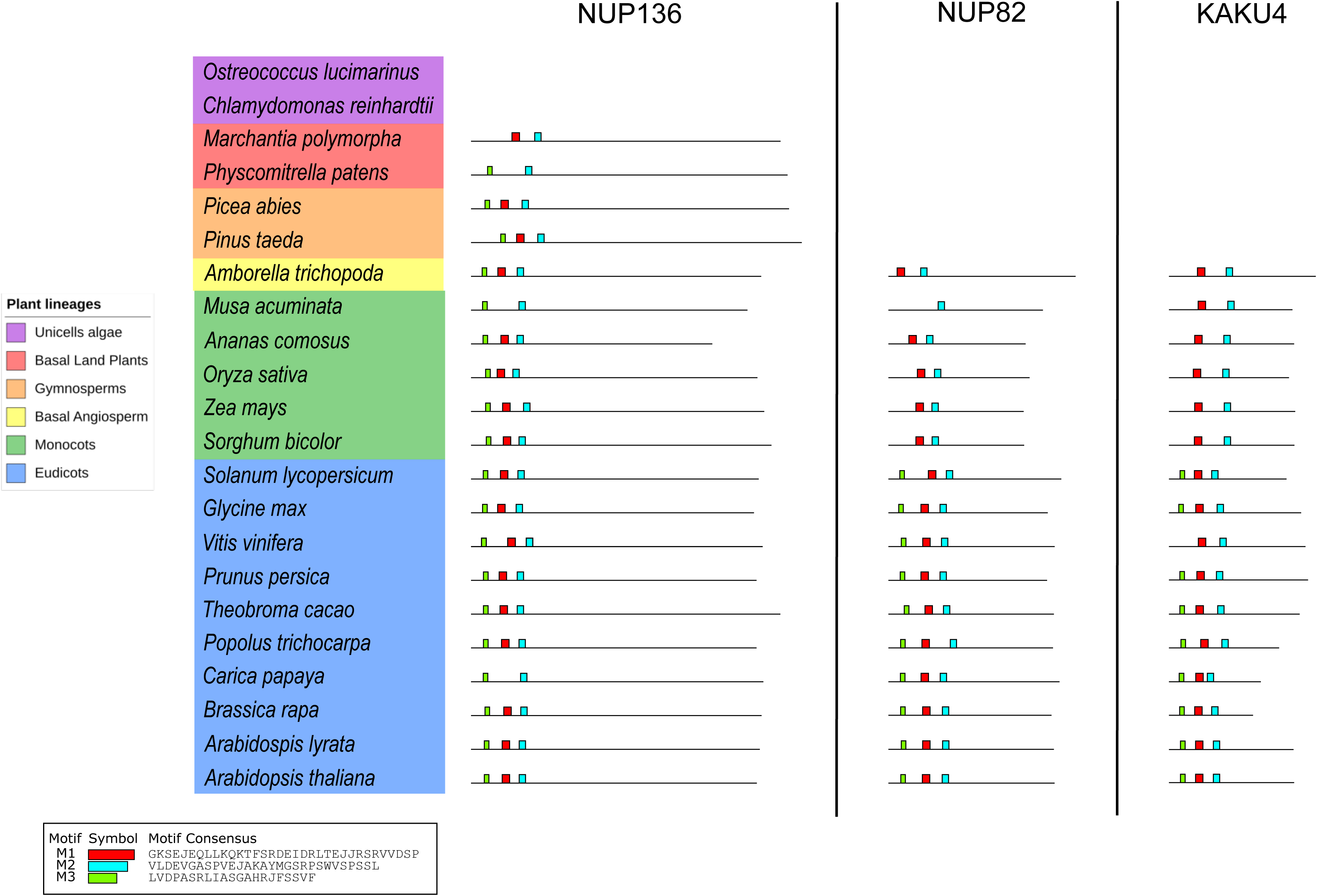
Summary of MEME results obtained with 52 orthologs from KAKU4, NUP82 and NUP136. **Left panel:** 22 plant species used in this study. From top to bottom: Unicellular algae (purple), Basal land plants (red), Gymnosperms (orange), Basal Angiosperms (yellow), Monocotyledons (green), and Dicotyledons (blue). **Right panel:** MEME suite results obtained with the 52 combined orthologs of NUP136, NUP82 and KAKU4. M3 (green), M1 (red) and M2 (light blue) sequences are indicated at the top. Accession numbers used in this study can be found in Supplementary table S2.

**Supplemental Figure S3:**
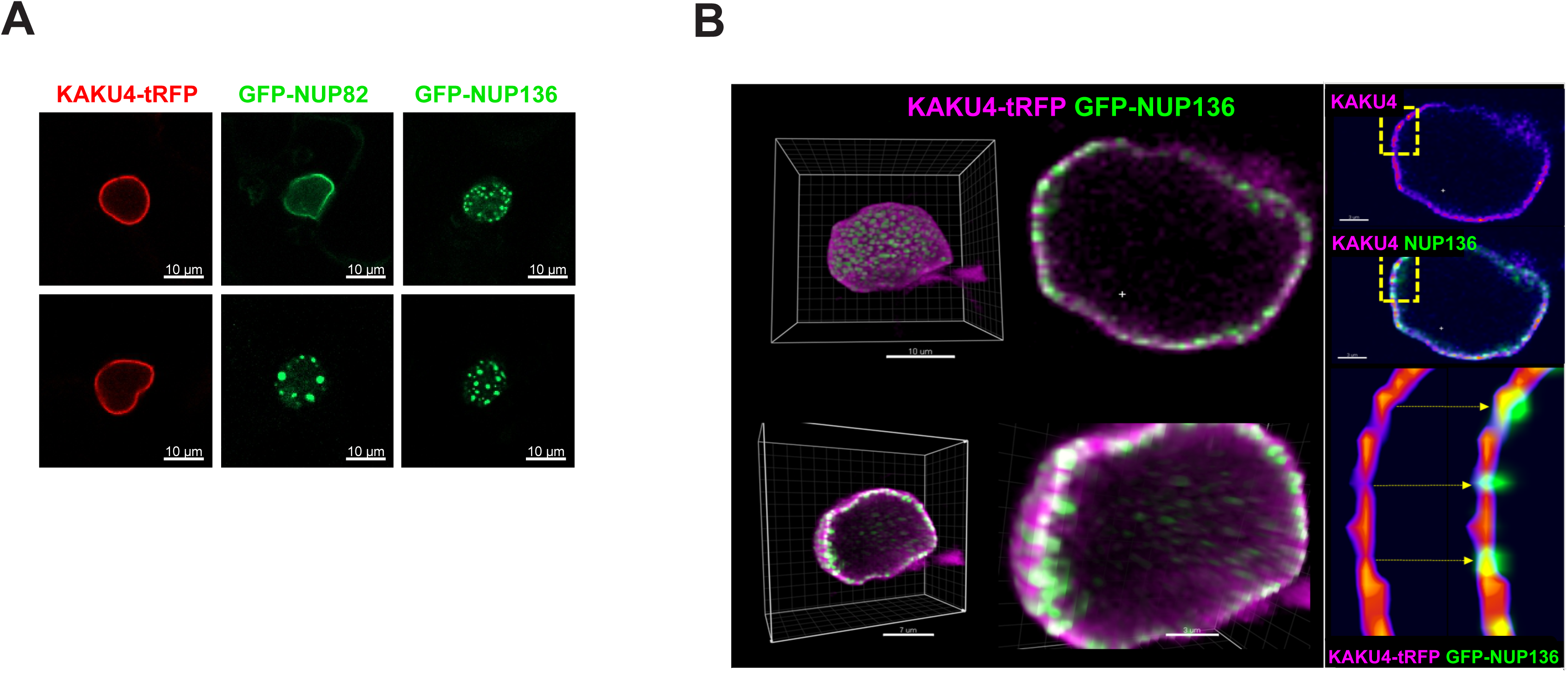
Localization of KAKU4, NUP82 and NUP136 at the nuclear periphery. **(A)** Localization of KAKU4-tRFP, GFP-NUP82-GFP and GFP-NUP136-GFP constructs in transient expression assay in *Nicotiana benthamiana* leaves. Two examples (top and bottom panels) are given. In a typical transient expression assay, KAKU4 localizes as a more uniform profile at the nuclear periphery (100% uniform profiles) than NUP82 that displays either a uniform pattern (57%) or large aggregates (43%). NUP136 always displays a spotty pattern (100%). **(B)** Co-expression between KAKU4 and NUP136 in transient expression assay in *Nicotiana benthamiana* leaves. Rendering of 3D view using the Imaris software showing the localization of KAKU4- tRFP between spots of NUP136-GFP expression. The right-bottom panel shows a zoom in a specific region showing alternating NUP136 (green spots) and KAKU4 (red segments between the green spots).

**Supplemental Figure S4:**
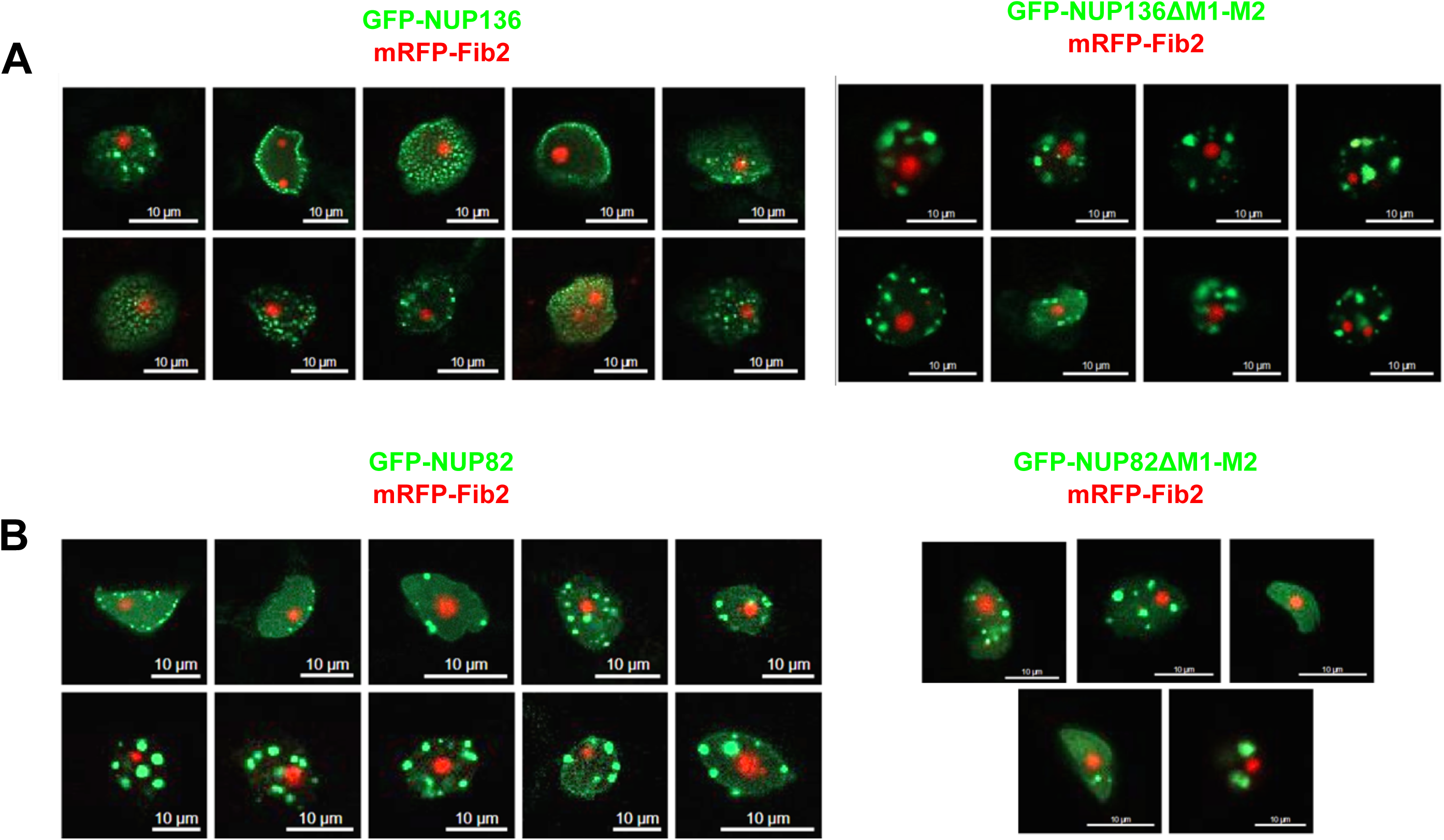
Impact of M1 and M2 motif deletions on *Arabidopsis thaliana* NUP82 and NUP136 localization. Confocal images of transiently expressed *N. benthamiana* leaves of **(A)** GFP-NUP136 and GFP-NUP136ΔM1-M2 (upper panel) and **(B)** GFP-NUP82 and GFP-NUP82ΔM1-M2 (lower panel) constructs co-transfected with mRFP-Fibrillarin.

**Supplemental Figure S5:**
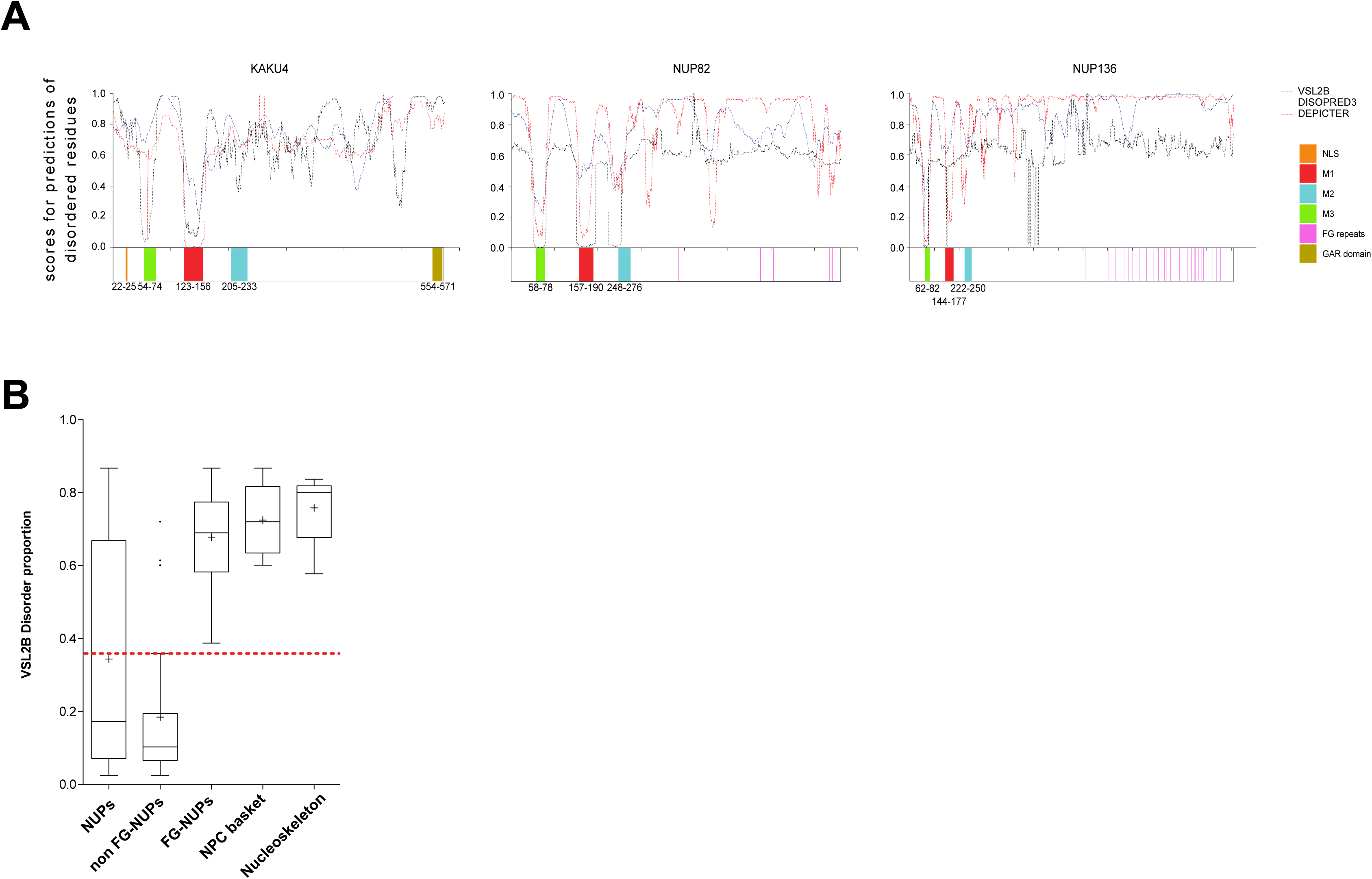
Predictions of intrinsically disordered regions in KAKU4, NUP136 and NUP82. **(A)** Profiles of intrinsically disordered residues in KAKU4, NUP136 and NUP82 proteins from *A. thaliana* predicted by VSL2B, DISOPRED3 and DEPICTER. A schematic representation of the three proteins is given below the disorder profiles with M3, M1 and M2 peptide motifs indicated in green, red and blue respectively. **(B)** VSL2B prediction from *A. thaliana* using DescribeProt precomputed predictions. From left to right: 31 NUPs, 21 non FG-NUPs, 10 FG-NUPs, 5 NUPs from the NPC basket and 5 nucleoskeleton proteins as described in **Supplemental Table S3**. Whiskers represent the Tukey intervals, Boxes represent interquartile distances, the horizontal line across whiskers represents the median, and “+” the mean values. The median value of proteome disorder from *A. thaliana* is indicated as a red dashed line.

**Supplemental Figure S6:**
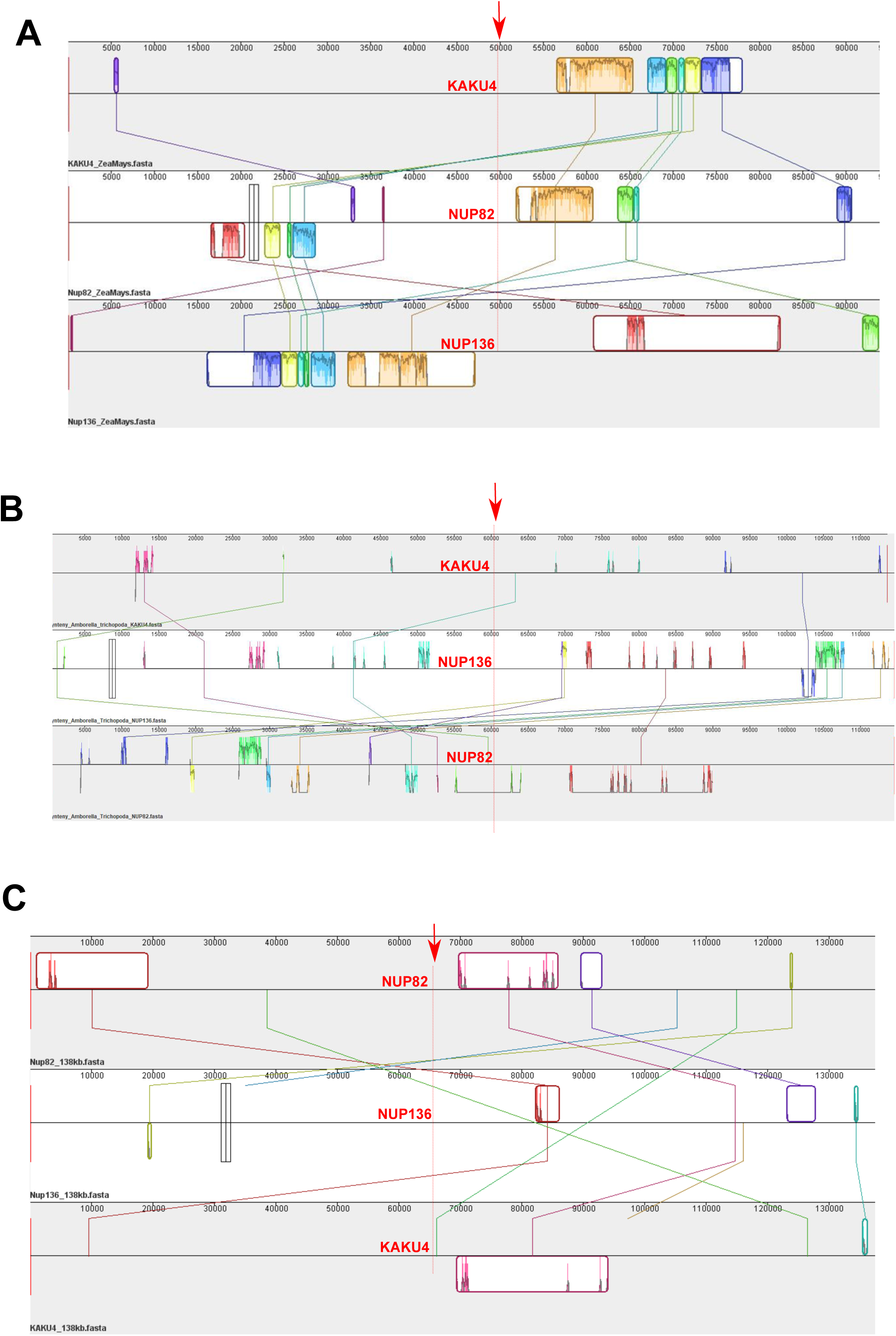
Locally conserved block and evolution rate of NUP136, NUP82 and KAKU4 in *Zea mays*. Locally conserved block (LCBs) between the genomic regions containing *KAKU4*, *NUP82* and *NUP136* orthologs in **(A)** *Zea mays* **(B)** *Amborella trichopoda* and **(C)** *Arabidopsis thaliana*. Graphical representation of LCBs detected by MAUVE (40) tool was performed for a genomic region of about 100kb each centered on the loci *KAKU4, NUP82* and *NUP136* (indicated by a red arrow). LCBs are indicated as contiguously colored region indicating the absence of genetic rearrangements of the chromosomal regions. Relationships of similarity between each region are highlighted by LCBs of the same color delimited by a frame and linked by lines of the same color. The similarity score is represented by peaks inside LCBs.

